# Design of a Conductive Hydrogel Coating to Improve Catheter–Tissue Coupling in Radiofrequency Ablation

**DOI:** 10.64898/2026.05.04.722766

**Authors:** Derek Bashe, Octavio Jalife, Archita Duvvada, Bhanaviya Venkat, Lukas Jaworski, Drew Bernard, Mathews John, Allison Post, Mehdi Razavi, Elizabeth Cosgriff-Hernandez

## Abstract

Radiofrequency ablation is a mainstay of cardiac rhythm management despite high recurrence rates. Current radiofrequency ablation catheters are limited by poor contact with trabeculated cardiac tissue that promotes uneven heating with hot spots that cause collateral damage and regions of incomplete ablation that promote recurrence. Herein, we report on a conductive hydrogel coating of radiofrequency ablation catheters to improve tissue contact while promoting efficient energy transfer. A method was developed to graft a polyether urethane diacrylamide hydrogel to the distal tip of the catheter that maintained stable adhesion following drying, sterilization, and rehydration. The coating also remained intact after passage through an introducer sheath and 50 cycles of radiofrequency ablation at clinical power. The hydrogel-coated catheter demonstrated enhanced tissue contact that was dependent on hydrogel modulus. Hydrogel-mediated ablation prevented steam pop incidence and generated homogeneous lesions in an ex vivo ablation model; however, increased hydrogel conductivity is needed to achieve comparable lesion dimensions as the bare metal catheter and prevent coating damage at higher power. Collectively, these results establish a tunable hydrogel coating method that addresses limitations of conventional radiofrequency ablation and offers a promising approach to enhance the safety and efficacy of cardiac ablation therapies.

## INTRODUCTION

Cardiac arrhythmias will affect one in every four Americans by age 40.^1–3^ Radiofrequency ablation is the mainstay for cardiac rhythm management, impacting 2.25 million patients each year with a $6B economic footprint.^4^ This ablative procedure blocks abnormal electrical signals that cause arrhythmia by destroying areas of electrical dysfunction. Despite its prevalence, current ablative techniques exhibit up to a 40% recurrence rate and can cause life-threatening complications such as atrio-esophageal fistula, steam pops, char microembolisms, and bleeding.^5–12^ Limitations of current radiofrequency catheters have been attributed to the inadequate contact between the rigid metal catheter tip and topographical cardiac tissue, leading to uneven heating with local hot spots that cause collateral damage or regions of incomplete ablation that promote recurrence.^13, 14^ Hypotonic irrigation, which enhances conductivity around the catheter tip, has been used to better control temperature and prevent char.^15^ However, it can lead to cell swelling, hemodilution, and electrolyte imbalance, and does not always improve lesion homogeneity or prevent gaps in lesions.^16^ Clinicians utilize increased contact force to improve tissue contact, but this can lead to dangerous complications, risking tissue perforation and atrio-esophageal fistula particularly in areas with thinner tissues.^17^ This method is also insufficient in improving tissue contact in areas with more irregular trabeculations in the atria.^18^ Overall, cardiac care lacks a solution for safely treating arrhythmia and effectively preventing recurrence, signifying the critical demand for innovation in ablation technology.

Given that poor conformation of current radiofrequency ablation catheter tips underpins the formation of hotspots and inadequate lesion formation, it follows that enhancing tissue coupling can improve patient outcomes. As stated above, this is due to the more homogeneous transfer of energy to 1) improve efficacy by preventing lesion gaps that lead to recurrence and 2) reduce complications by preventing ‘hot spots’ that lead to tissue damage. We hypothesize that a soft, conductive interface for the ablation catheter would enhance tissue coupling and provide the means to improve safety and efficacy without increasing contact force. Conductive hydrogels are promising for this application due to their soft tissue-like mechanics, which allow improved conformation to the irregular topology of the endocardium and ability to transduce radiofrequency energy. Utilizing slabs of an ionically conductive hydrogel atop myocardial tissue, we previously demonstrated that hydrogel-mediated radiofrequency ablation results in more homogeneous lesion formation at standard ablation power and duration while eliminating steam pops.^19^ In a tissue stack model of atrial and esophageal tissue, hydrogel-mediated ablation further protected tissues from thermal damage by preserving underlying esophageal tissue while maintaining consistent lesion geometry. In contrast, uncoated radiofrequency ablation probes caused significant tissue damage, steam pops, and cavitation. To translate this finding into an off-the-shelf device suitable for clinical use, a method is needed that can apply this conductive hydrogel as a coating of radiofrequency ablation catheters to serve as an interface between the ablation electrode and the endocardial tissue. Critical to the development of this technology is the use of grafting methodologies to ensure conformal contact with atrial tissue and mechanical and electrical durability throughout deployment.

In this work, we report on the development of a hydrogel coating for ablation catheters and investigated the coating’s ability to improve tissue coupling, and thus energy transfer, while maintaining physical integrity in simulated clinical use. First, hydrogel coatings were fabricated, dried, and sterilized with ethylene oxide to establish an appropriate sterilization protocol. To ensure clinical utility, the coated catheters were tested for both mechanical and electrical durability. Damage resistance was assessed following equilibrium swelling in 0.9% saline, passing through an introducer sheath, and enduring multiple cycles of radiofrequency ablation at clinical power. A model of atrial tissue was utilized to establish the relationship between coating stiffness and thickness and the resulting tissue contact area. Hydrogel stiffnesses were selected to bracket atrial tissue stiffness, encompassing values below, matching, and above native atrial myocardium. Finally, hydrogel-mediated ablation was carried out in an ex vivo model to investigate lesion formation and steam pop occurrence compared to the traditional bare metal catheter. Collectively, this work establishes a durable hydrogel coating method for radiofrequency catheters and demonstrates the potential hydrogel-mediated ablation to improve ablation procedures.

## RESULTS AND DISCUSSION

Our lab previously synthesized an ionic hydrogel based on polyether urethane diacrylamide (PEUDAm) that demonstrates enhanced durability, tunable stiffness, and suitable conductivity (12.8 ± 1.5 mS/cm) for ablation applications.^20, 21^ In this work, we developed a coating methodology to apply this hydrogel to radiofrequency ablation catheters. The metal catheter tip was first cleaned thoroughly and dried prior to treatment with oxygen plasma to introduce hydroxyl groups (**Figure 1A,B**). The surface was then functionalized with a bifunctional organosilane coupling agent, 3-(trimethoxysilyl) propyl methacrylate (TMSPMA), to enable grafting of the PEUDAm to the surface.^22^ A series of TMSPMA concentrations in ethanol were tested to determine a concentration that provided stable grafting. Ethanol was selected as the solvent to facilitate hydrolysis and condensation. A 10% TMSPMA concentration was found to provide controlled silane layer formation and prevent total self-polymerization of TMSPMA. After one hour submersion in the TMSPMA solution, the catheter tip was rinsed in ethanol and dried under a slow, steady flow of nitrogen that allowed for uniform evaporation of ethanol for an even layer of TMSPMA without chemically interacting with the silane layer (**Figure 1C**). The hydrogel precursor solution was then loaded into a cylindrical mold with dimensions chosen to match the shape of the catheter tip, allowing for a uniform coating. The mold was rinsed in Sigmacote prior to adding the hydrogel precursor solution and inserting the catheter tip. An ultraviolet exposure duration of 4 minutes was identified to initiate radical formation and promote uniform crosslinking of the hydrogel onto the catheter tip. After photocuring, the catheter tip was carefully removed from the mold and the hydrogel coating was soaked in deionized water to remove unreacted reagents (**Figure 1D**).

**Figure 1.**
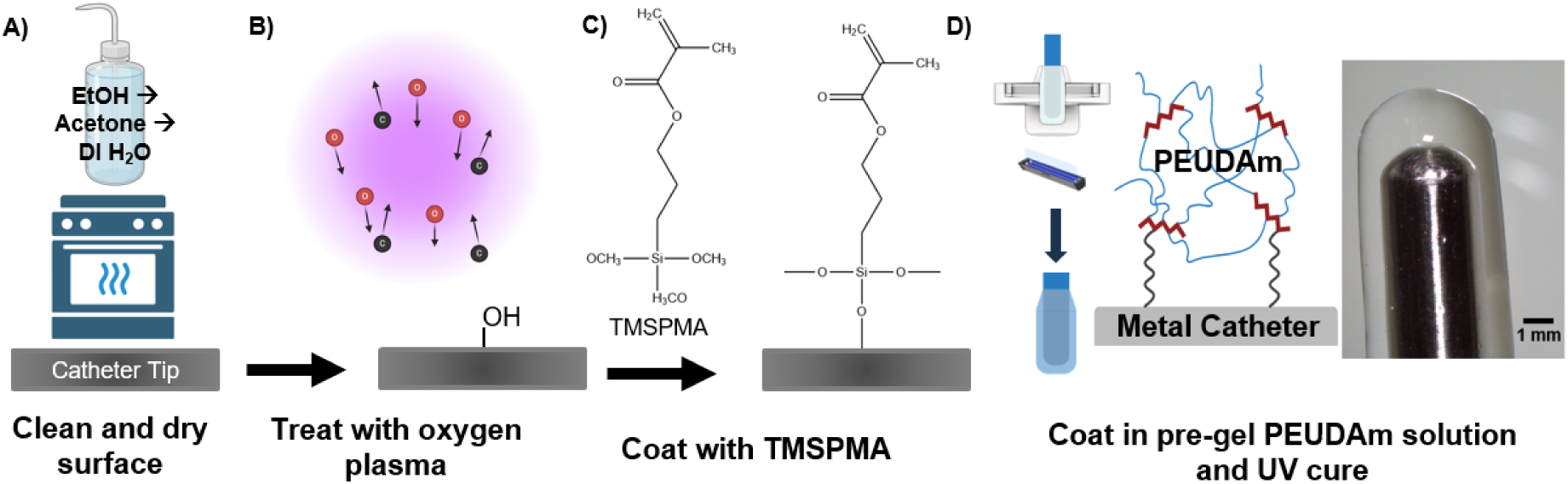
Fabrication of the ablation catheter coating. The hydrogel coating was covalently crosslinked directly onto the catheter tip using TMSPMA functionalization. a) First, the catheter tip was cleaned with ethanol, acetone, and DI water and then dried for one hour. b) Oxygen plasma was used to modify the metal’s surface with hydroxyl groups. c) The catheter tip was then soaked in a 10% TMSPMA in ethanol for 1 hour. d) Finally, the catheter was removed from the TMSPMA solution, rinsed in ethanol, dried with nitrogen gas, and submerged in the hydrogel precursor solution using a tapered mold prior to curing with UV light (365nm) for 4 minutes.

### Hydrogel Coating Stability

Following successful development of the coating methodology, off-the-shelf potential and durability throughout a simulated ablation procedure was tested. We envision that the single-use hydrogel-coated ablation catheter will be stored dry after sterilization and then rehydrated in the surgical suite prior to deployment. Therefore, the coating must resist swelling-induced delamination following drying, sterilization, and rehydration to be viable as an off-the-shelf device. The effect of ethylene oxide sterilization on the properties of a similar PEG-based hydrogel was previously tested and reported to have minimal effect on hydrogel properties or the polymer backbone itself as examined via NMR analysis.^23–30^ Similarly, the hydrogel coating remained intact following drying, sterilization and rehydration with no notable difference in swelling or dimensions (**Figure 2A**).

**Figure 2.**
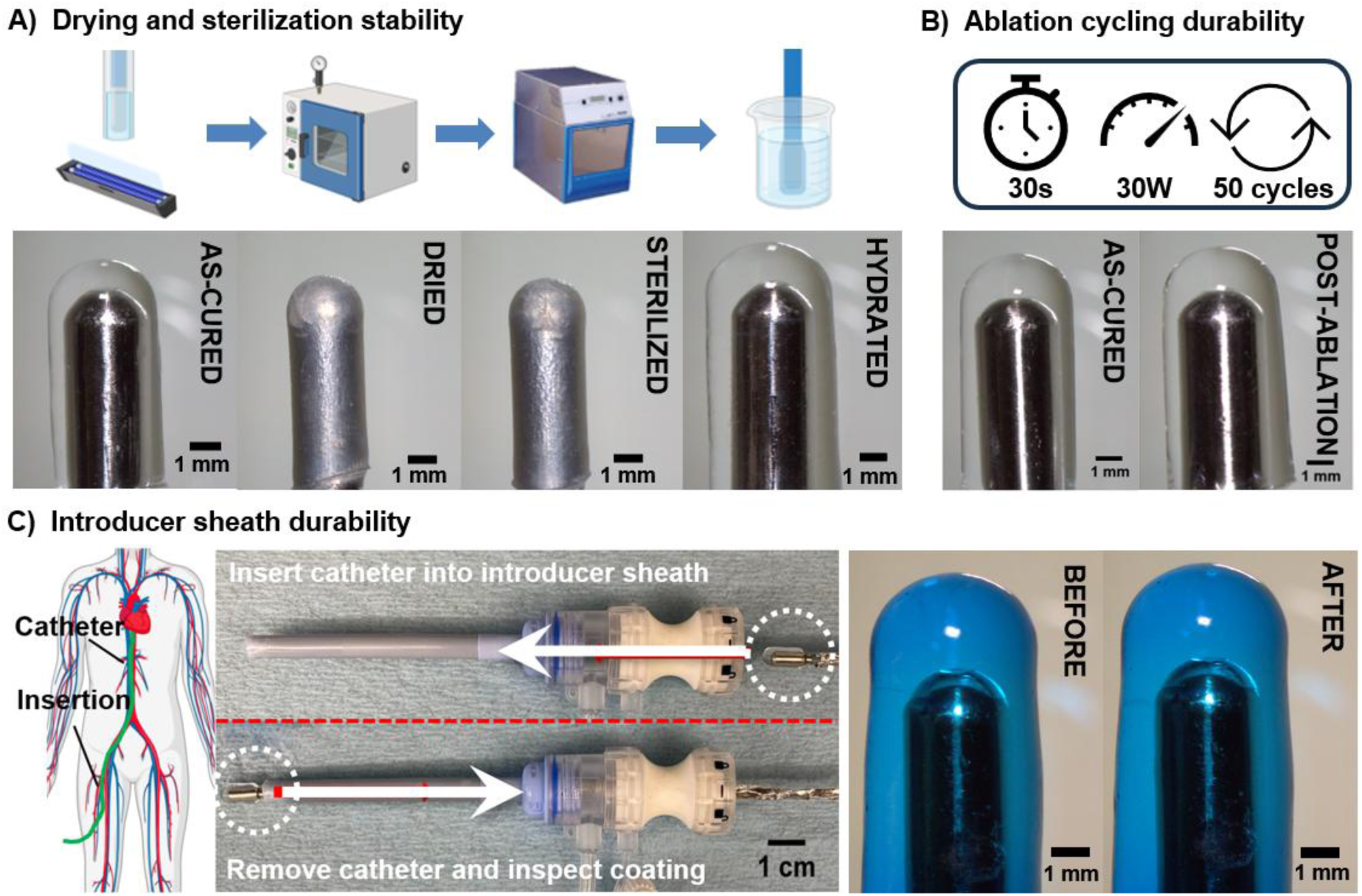
Hydrogel coating durability. Stereoscope images of 20% 10kDa PEUDAm catheter coating before and after a) drying, Ethylene Oxide (EtO) sterilization, and swelling to equilibrium in DI water, b) 50 cycles of radiofrequency ablation at 30W, and c) introducer sheath deployment. n = 3 for each composition.

The coatings must also withstand the ablation procedure which can include 10s of ablation cycles (30s on 30s off). The electrical durability of the hydrogel-coated ablation catheter was investigated using a radiofrequency generator and submerging the catheter in a tank filled with saline at 37 °C. There was no apparent damage or delamination of the hydrogel coating after 50 cycles at 30 W (**Figure 2B**). Mechanical durability is also a key criterion in the deployment of the ablation catheter as damage may induce particulate embolization or full delamination, ultimately eliminating the coating’s functional advantages. In particular, passing in and out of the introducer sheath represents a critical step in the procedure with respect to potential hydrogel damage or delamination. In cardiac ablation, 6 to 14 Fr sheaths are typically inserted via the femoral vein and incorporate a saline-inflated valve to prevent backflow of blood.^31^ While metal catheter tips can easily traverse this valve, this interface presents a potential site of high shear stress on the hydrogel coating. Therefore, we characterized coating damage resistance with an introducer sheath test, in which the coating was inserted into and pulled back through the valve of an introducer sheath. Notably, the coating remained intact with no observable damage or delamination (**Figure 2C**).

### Effect of Hydrogel Coating on Tissue Contact

Tissue contact serves as the primary target for improving radiofrequency ablation, as it is a key determinant of improved outcomes for patients undergoing cardiac ablation procedures. A custom model of atrial tissue matching physiological stiffness was developed and utilized to estimate the effect of coating thickness and stiffness on the area of contact between the catheter and atrial tissue. First, we examined the effect of hydrogel coating thickness on tissue contact area. Two mold dimensions (D = 4 mm and 6 mm) were selected based on atrial trabeculation geometry and introducer sheath constraints. The hydrogel stiffness was kept by using a single composition of 10% 20kDa PEUDAm. As expected, there was a small increase in hydrogel-coated probe diameter after equilibrium swelling. Increased mold diameter resulted in increased coating thickness and an associated increase in contact area with the atrial tissue mimic at standard contact force of 20 g (**Figure 3**).

**Figure 3.**
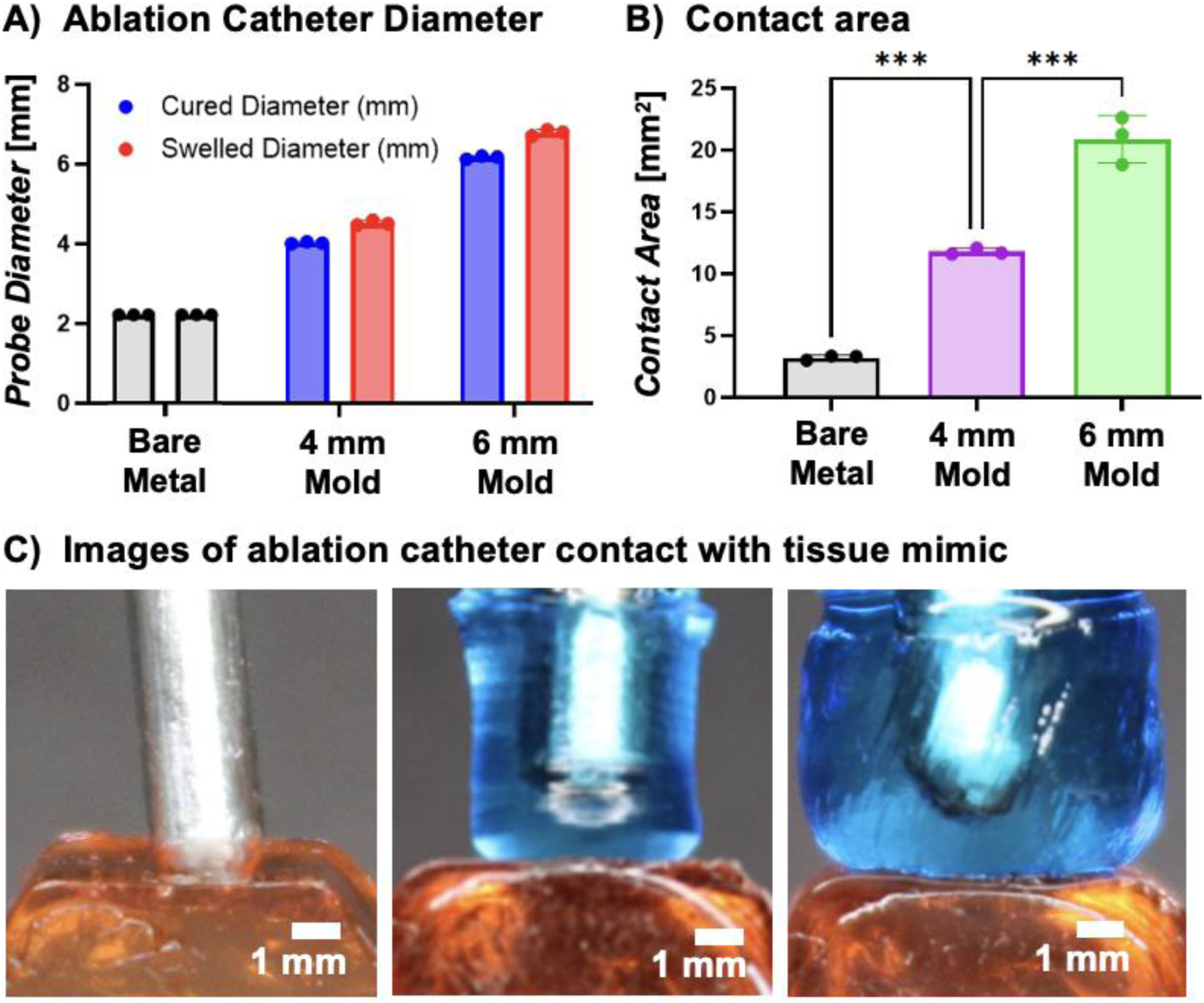
Effect of hydrogel coating thickness on contact area. a) Catheter diameter before and after swelling to equilibrium; b) contact area between ablation catheter and atrial tissue mimic, as determined by arc length analysis; c) images of ablation catheter contact with atrial tissue, 20 g contact force. Data for all comparisons represent individual data points, means, and standard deviations of each specimen (n = 3 for each composition, 3 compositions). Significant differences between groups are marked as follows: ∗∗∗ p < 0.0002.

Next, we investigated the effect of hydrogel coating stiffness on contact area while holding the mold diameter constant at 4 mm. Hydrogel compositions were identified with stiffness spanning below to above the physiological stiffness ranges of atrial tissue, which lies between 100-250 kPa (**Figure 4A**).^32^ We hypothesized that if coating stiffness is less than that of the atrial tissue, the coating will deform to the surface of the tissue, increasing contact area to promote uniform energy delivery. Conversely, if coating modulus is greater than that of the tissue, the tissue will deform first, increasing procedural risk such as atrio-esophageal fistula.^17^ Bulk hydrogel stiffness for each composition was evaluated to confirm our chosen compositions bracketed atrial tissue stiffness, **Figure 4A**.^32^ The 10% 20kDa PEUDAm hydrogel exhibited moduli below those of atrial tissue (71.9 ± 4.5 kPa), the 20% 10kDa PEUDAm hydrogel fell within the physiological range (195.0 ± 21.7 kPa), and the 30% 3.4kDa PEUDAm hydrogel exceeded it (530.0 ± 90.6 kPa). All compositions were successfully fabricated as ablation catheter coatings and increased contact area as compared to the bare-metal ablation catheter (**Figure 4B-D**). There was a marked increase in contact area with decreasing hydrogel coating stiffness that was attributed to greater deformation to the atrial tissue mimic for improved contact. All coated catheters have similar diameters; therefore, the significant increase in contact area with decreasing coating stiffness supports the hypothesis that softer coatings undergo greater deformation. The 30% 3.4kDa PEUDAm coating showed minimal deformation and did not substantially increase contact area beyond what would be expected from its larger diameter relative to the bare metal catheter. In contrast, the 20% 10 kDa and 10% 20 kDa PEUDAm coatings produced significantly greater contact areas than can be explained by probe thickness alone under standard contact force, indicating marked hydrogel deformation that increased contact area.

**Figure 4.**
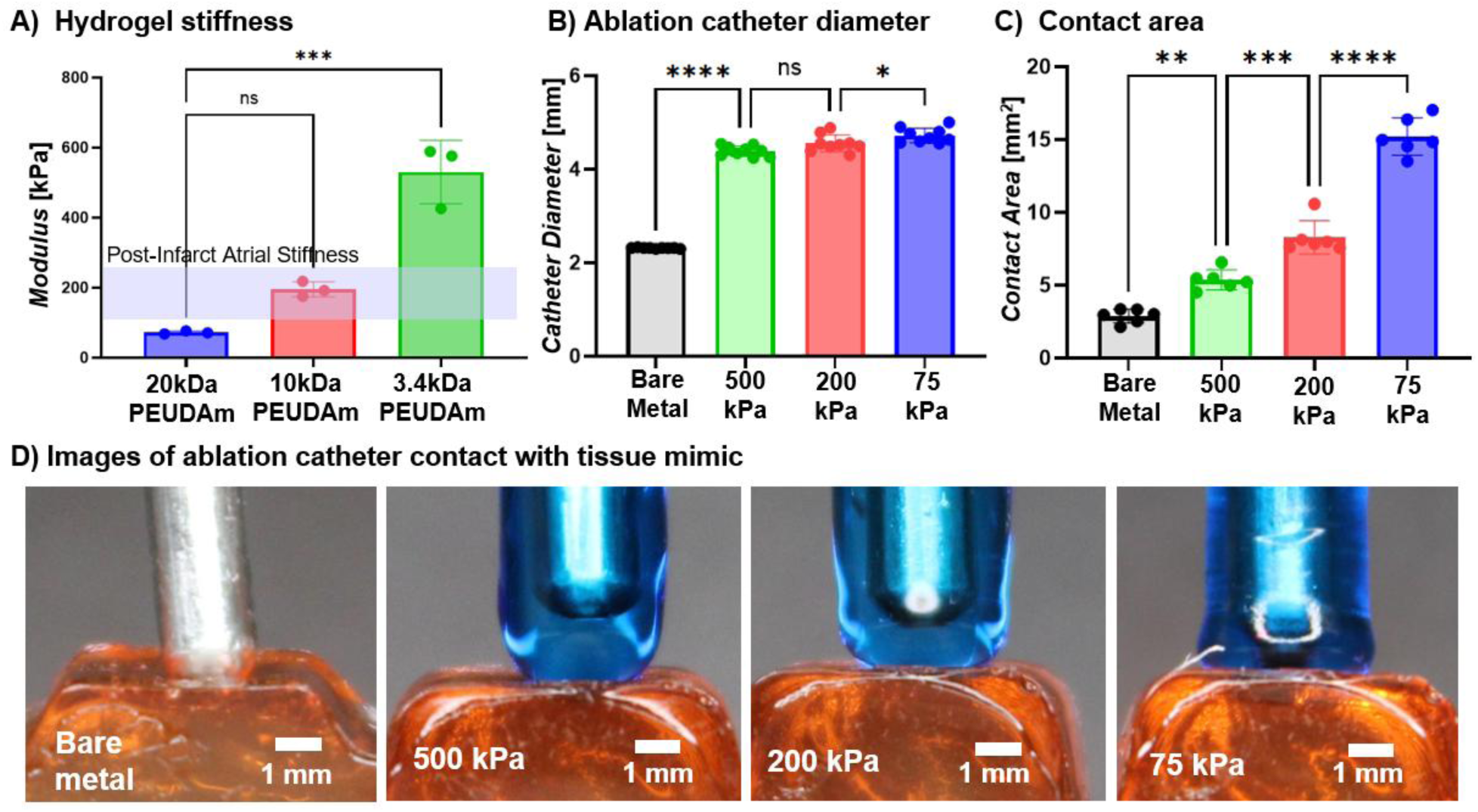
Effect of hydrogel coating stiffness on tissue coupling. Hydrogel coatings with compositions of 10% 20kDa PEUDAm, 20% 10kDa PEUDAm, and 30% 3.4kDa PEUDAm. a) hydrogel stiffness in comparison to atrial stiffness (n = 3 for each composition, 3 compositions); b) probe diameter across hydrogel compositions and bare metal (n = 9); c) contact area between ablation catheter and atrial tissue mimic, as determined by arc length analysis; d) images of ablation catheter contact with atrial tissue, 20 g contact force. Data for all comparisons represent individual data points, means, and standard deviations of each specimen (n = 6 for each composition, 3 compositions). Significant differences between groups are marked as follows: ∗ (p < 0.05), ∗∗ (p < 0.001), ∗∗∗ (p < 0.0002), ∗∗∗∗ (p < 0.0001).

To investigate coating durability across these compositions, physical integrity after drying and sterilization was again assessed. The softer coatings (10% 20kDa and 20% 10kDa PEUDAm) displayed no damage upon rehydration; whereas the stiffer coating composition (30% 3.4kDa PEUDAm) cracked during the drying phase and delaminated from the metal catheter tip (**Figure S1**). This is likely a result of its elevated stiffness limiting the ability of the network to accommodate shrinkage. Subsequent swelling induced delamination, consistent with failure to maintain interfacial adhesion at the TMSPMA-functionalized metal surface. Electrical durability was again investigated with ablation cycling; all coating compositions successfully maintained physical integrity throughout this testing (**Figure S2**). Finally, introducer sheath durability testing was repeated. None of the coating compositions experienced delamination; however, stiffer coatings were more reliable in that no particulates were generated, while some lower-modulus coatings exhibited minor surface damage (**Figure S3**). Although reduced modulus may improve tissue contact, coating integrity is essential for safe deployment, as failure could result in loss of coating function or, more severely, embolic risk from generated particulates.^33^ This limitation may be addressed through the taper design of the acrylic molds, and further mitigated by optimization of introducer sheath selection to reduce insertion- or removal-induced mechanical stress. As coating composition undergoes further iteration, mechanical durability will continue to be assessed and improvements in mold geometry and thickness will be identified to achieve a balance of durability and tissue conformation.

### Effect of Hydrogel-Mediated Ablation on Lesion Geometry and Tissue Damage

Lesion formation was characterized using an ex vivo model for each coating composition in comparison to the bare metal catheter. Lesions of porcine ventricular free wall tissue were compared after 30-second radiofrequency ablations at 30W (**Figure 5A**). All hydrogel-coated ablation catheters generated even lesions with lesion depth and width increasing with decreasing hydrogel coating stiffness. This corresponded to increasing contact area; however, the comparison to the bare metal ablation catheter did not show a similar correlation to contact area. Lesion dimensions were statistically similar or greater for the bare metal ablation catheter as compared to the hydrogel-coated catheters, **Figure 5B, C**. We hypothesize that the reduced ablation lesion size was due to reduced current density and enhances heat dissipation in hydrogel-mediated ablation.

**Figure 5.**
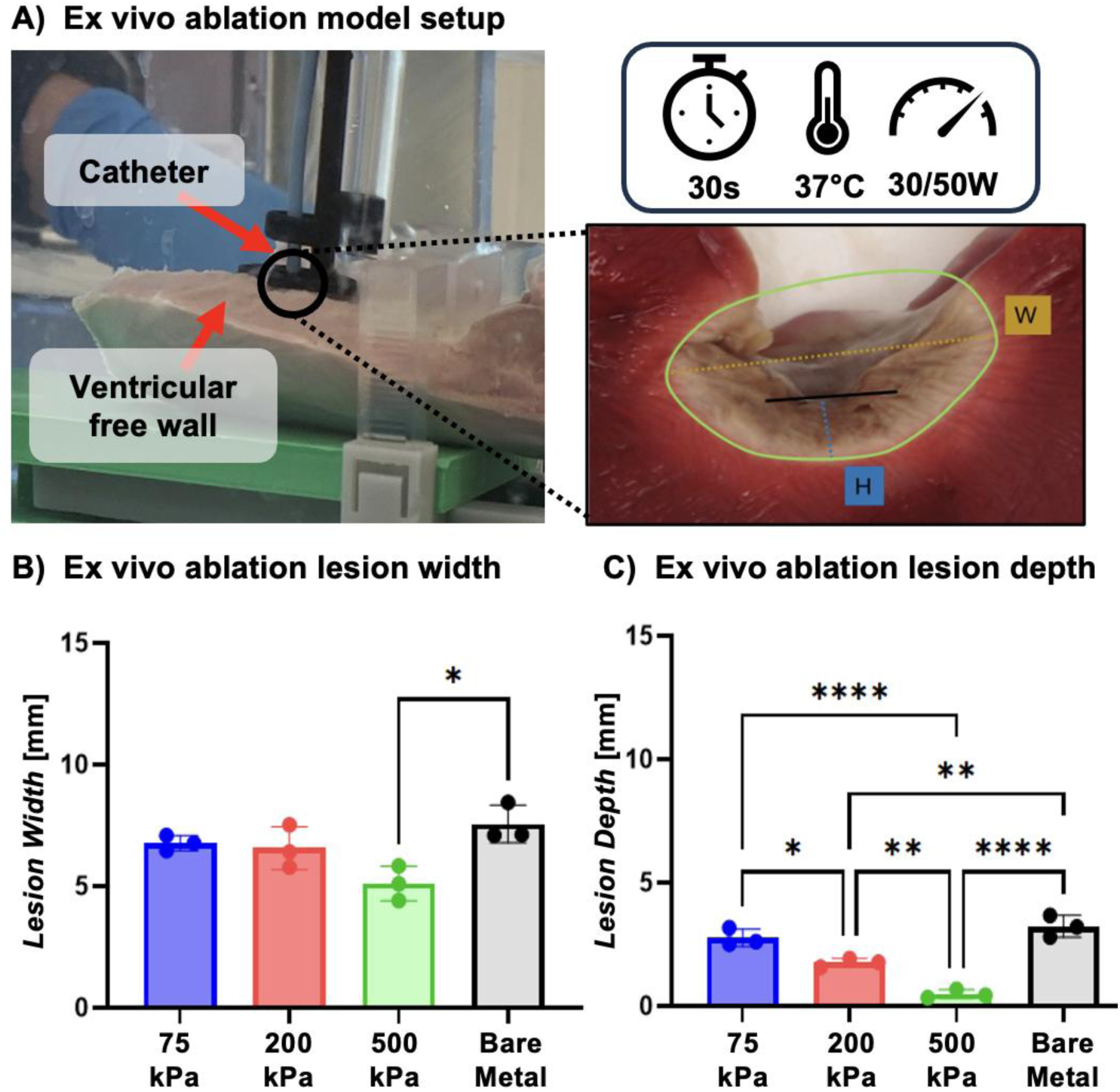
Effect of hydrogel-mediated ablation on lesion geometry at 30W. a) Setup to carry out ex vivo ablation on ventricular free wall. Nonviable tissue presents as light brown while viable tissue is stained bright red. b) Ex vivo lesion widths compared to a standard ablation catheter at 30W. e) Ex vivo lesion depths compared to a standard ablation catheter at 30W. Data for all comparisons represent individual data points, means, and standard deviations of each specimen (n = 3 for each composition, 3 compositions). Significant differences between groups are marked as follows: ∗ (p < 0.05), ∗∗ (p < 0.001), ∗∗∗ (p < 0.0002), ∗∗∗∗ (p < 0.0001), ns (p > 0.05).

Ablation power is another variable used to control lesion size clinically. At an increased power of 50 W, all hydrogel compositions successfully prevented steam pops. Hydrogel-mediated lesions were qualitatively more homogeneous than the lesions formed by a standard ablation catheter, and fell within the range of lesion geometries required to prevent recurrence of arrhythmia.^34^ Bare metal stents exhibited an average 2.88 ± 0.47 steam pops per 30 seconds with a corollary tissue cavitation (**Figure 6A**). Lesion depths and widths were markedly greater at 50 W for the bare metal probe (**Figure 6B, C**). There was a similar increase in lesion size at higher power for hydrogel-mediated ablation but not at the magnitude observed with bare metal. Notably, the stiffest hydrogel coating (30% 3.4 kDa PEUDAm) formed more shallow lesions at 30 W and were unable to form lesions at 50 W due to a maximum-impedance safety mechanism being activated. Radiofrequency ablation relies on Joule (resistive) heating, wherein tissue heating occurs as the radiofrequency energy encounters resistance during propagation through tissue.^35^ A similar mechanism may arise within the hydrogel coating if it is excessively thick or insufficiently conductive. We hypothesized that reduced energy transfer to the tissue due to increased impedance may reduce lesion size. Hydrogel resistive heating may lead to microbubble formation and partial delamination of the hydrogel coating which would substantially increase the impendence and prevent lesion formation.

**Figure 6.**
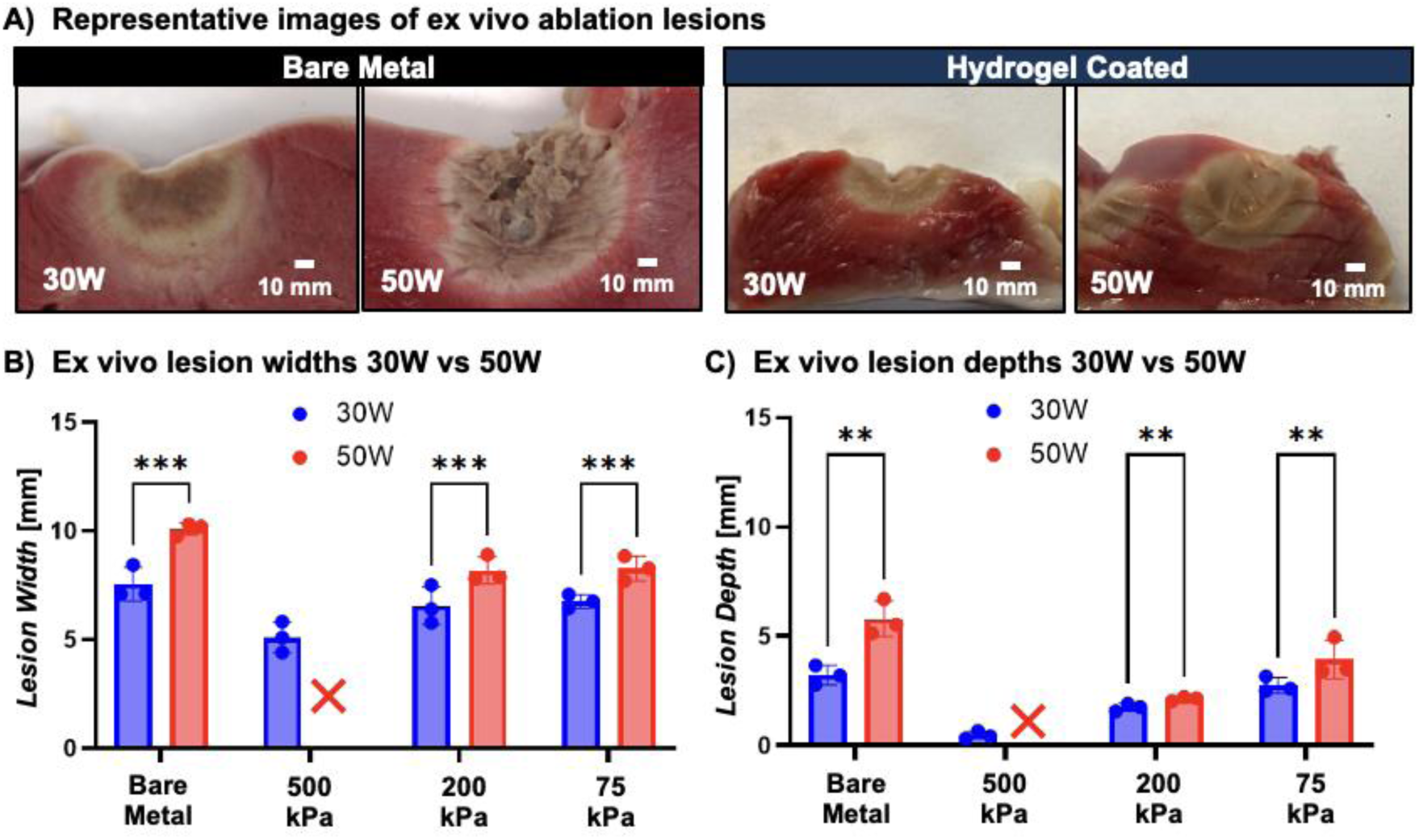
Effect of hydrogel-mediated ablation on lesion geometry and safety at increased ablation power. a) Representative lesions formed by bare metal and hydrogel-coated catheters at 50W, demonstrating increase in lesion dimensions at increased power and tissue cavitation due to steam pop for the bare metal catheter ablation (scale bar is 10 mm). b) comparison of ablation power (30W vs 50W) on ex vivo lesion width. c) comparison of ablation power (30W vs 50W) on ex vivo lesion depth. Data for all comparisons represent individual data points, means, and standard deviations of each specimen (n = 3 for each composition, 3 compositions). Significant differences between groups are marked as follows: ∗∗ (p < 0.001), ∗∗∗ (p < 0.0002). X indicates no lesion formation.

### Effect of Hydrogel Conductivity on Coating Stability at Higher Power

Previously, all coating compositions survived 50 cycles of ablation at a clinical power of 30 W, demonstrating their ability to match the procedural capabilities of a standard bare metal catheter. Increasing the power to 50 W resulted in hydrogel coating delamination (**Figure 7A**). We hypothesize that this delamination was due to resistive heating within the hydrogel that increased swelling and generated microbubbles to cause delamination. This is consistent with the reduced lesion size observed in the previous study. Increased ablation power is desirable because it will improve tunability of the ablation procedure itself. In theory, increasing the hydrogel conductivity will reduce resistive heating and allow for more efficient energy coupling with the tissue and prevent hydrogel coating damage. To test this hypothesis, we increased coating conductivity by increasing the concentration of saline from 0.9% to 2%, resulting in a 50% increase in conductivity. When coating conductivity was increased from 10 mS/cm to 15 mS/cm, all coatings remained intact with no evidence of delamination for the full 50 cycles at 50W power (**Figure 7B**). However, in a clinical situation this is not feasible; PEUDAm is inherently non-ionic, so the coating would equilibrate back to the conductivity of the extracellular environment surrounding the heart.^20^ Thus, future work will consist of maximizing coating conductivity to suppress resistive heating and enable stable, power-dependent lesion modulation. One potential strategy is to incorporate ionic species into the backbone of our polymer (e.g.; 2-Acrylamido-2-methylpropane sulfonic acid sodium salt).^21^ A key criterion in the design of a more conductive hydrogel coating will be to balance conductivity with mechanical toughness, requiring future reiteration of coating methodology as well as reexamination of coating durability throughout deployment.

**Figure 7.**
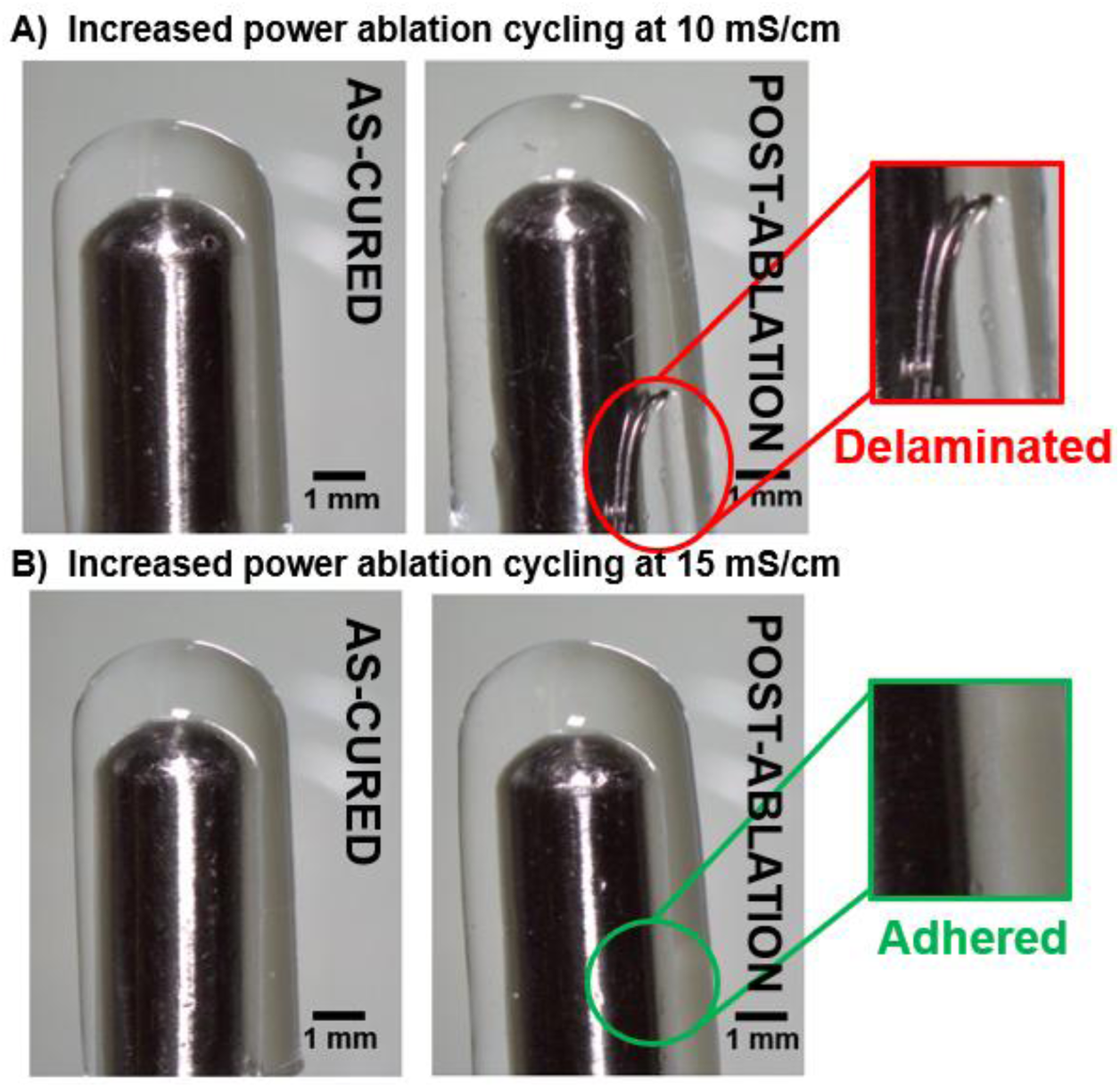
Effect of hydrogel conductivity on coating durability at higher ablation power. Stereoscope images of hydrogel-coated catheters before and after 50 cycles of 50W ablation with a) 20% 10kDa PEUDAm coating swelled in 0.9% saline with a hydrogel conductivity of ∼10 mS/cm and b) 20% 10kDa PEUDAm coating swelled in 2.0% saline with a hydrogel conductivity of ∼15 mS/cm.

### Development of a Tissue Mimic Model to Test Lesion Formation

To better isolate the role of heat permeation on lesion geometry and increase iterative testing, we developed a tissue mimic that matches atrial physiological stiffness and conductivity and incorporates a thermochromic dye that turns from black to white at 60 °C (**Figure 8A**). We then compared ablation lesion formation in this tissue mimic to the ex vivo model. This study demonstrated lesion width in the thermochromic model were consistent with those observed in ex vivo tissue at 30 W but with reduced variability (**Figure 8B**); however, lesion depths were statistically greater on the thermochromic model (**Figure 8C**). This is likely due to the fact that the surface of the thermochromic tissue mimic was completely flat, compared to the excised tissue which had topographical features. This model provides a reproducible and predictive model for iterative testing of ablation parameters and also isolate the effect of heat transfer from tissue variability.

**Figure 8.**
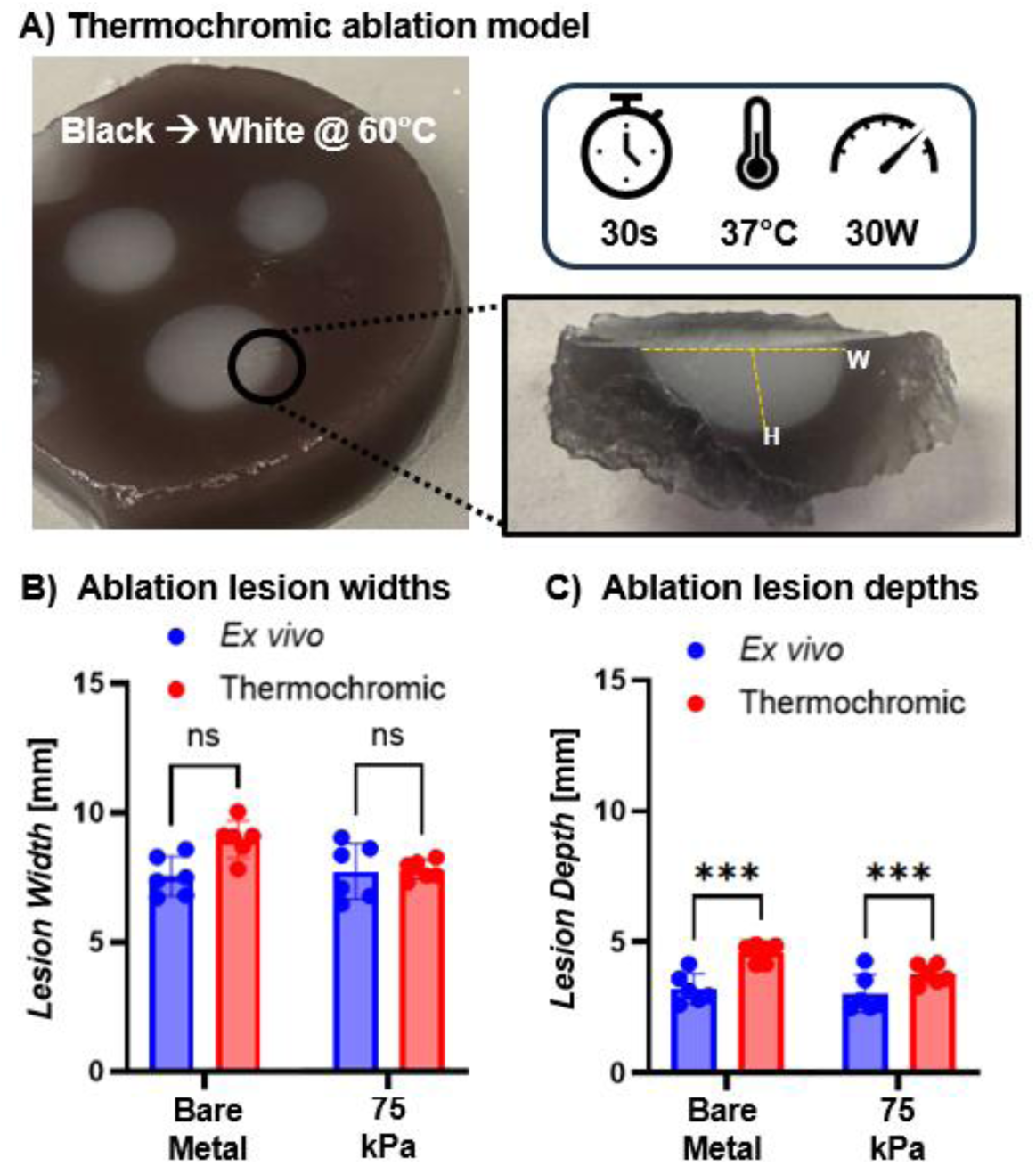
Thermochromic atrial tissue mimic as an analog for ex vivo ablation. a) Images of tissue mimic with “nonviable” tissue turning white after ablation and “viable” tissue remaining black; b) comparison of ex vivo vs thermochromic tissue mimic lesion width; c) comparison of ex vivo vs thermochromic tissue mimic lesion depth. Data for all comparisons represent individual data points, means, and standard deviations of each specimen (n = 6). Significant differences between groups are marked as ∗∗∗ p < 0.0001, ns indicates no significant differences, p > 0.05.

## CONCLUSIONS

In this work, we developed methodology to apply a hydrogel coating to radiofrequency ablation catheters and evaluated the effect on lesion formation and uniformity. By demonstrating that the hydrogel coating can withstand drying, sterilization, simulated deployment and ablation cycling, we confirmed the potential use of this device as an off-the-shelf ablation catheter in the clinic. The deformable and conductive properties of the coating provided a platform to improve tissue contact and homogeneous energy delivery, generating the foundation to reduce procedural risks and recurrence of arrhythmia. Specifically, the hydrogel coating showed increased contact area with atrial tissue due to an increase in deformation at lower stiffness as well as an increase in thickness at higher mold diameters, supporting the primary claim that these coatings can improve tissue contact. Lesion formation in an ex vivo model demonstrated the ability of the hydrogel coatings to form clinically-sized lesions that were more homogeneous than those formed via the bare metal ablation catheter while preventing damage due to steam pops. However, increased hydrogel conductivity is needed to achieve comparable lesion dimensions as the bare metal catheter and prevent coating damage at higher power. Overall, these studies established key design rules to guide the development and optimization of hydrogel-coated catheters and highlight the potential of these soft, conductive hydrogel coatings to improve safety and efficacy of cardiac ablation. In the future, we will expand the hydrogel design space through systematic variation in composition and mold geometries, guided by the above findings, to optimize tissue contact, conductivity, and durability throughout cardiac ablation.

## METHODS

### Materials

All reagents were purchased from Sigma-Aldrich and used as received unless otherwise noted.

### Synthesis of polyether urethane diacrylamide (PEUDAm)

PEUDAm was synthesized as previously reported.^36^ Briefly, PEG (3.4kDa, 10kDa, 20 kDa) was dissolved in DCM and reacted with CDI (1:15 molar ratio) under nitrogen for 2 h at 30-35 °C. The reaction continued mixing overnight prior to quenching with water and solvent removal to collect PEG-CDI. The product was washed with deionized water in a separatory flask, dried with sodium sulfate, and collected via vacuum filtration prior to precipitation in 10X volume of ice-cold ether and a subsequent vacuum filtration. Solvent was removed by drying under vacuum for 1 hour. PEG-CDI was added dropwise to EDA (1:15 molar ratio) under nitrogen and stirred for 24 h at room temperature. The resulting PEG-EDA was washed with deionized water in a separatory flask, dried with sodium sulfate, and collected via vacuum filtration prior to precipitation in 10X volume of ice-cold ether and a subsequent vacuum filtration. Solvent was removed by drying under vacuum for 1 hour. PEUDAm was then synthesized by functionalizing the PEG-EDA with acrylamide groups through the addition of triethylamine (1:4 molar ratio) and acryloyl chloride (1:4 molar ratio) to PEG-EDA. The reaction was stirred for 24 h then quenched using potassium carbonate. The resulting PEUDAm was washed with deionized water in a separatory flask, dried with sodium sulfate, and collected via vacuum filtration prior to precipitation in 10X volume of ice-cold ether and a subsequent vacuum filtration. The final product was dried at atmospheric pressure overnight, then briefly under vacuum. The chemical structure of the final PEUDAm product was verified with proton nuclear magnetic resonance H^1^ NMR (CDCl_3_): *δ* = 6.94 (broad s, 2H, –C–N*H*–CH_2_–), 6.28 (dd, 2H, *H*_2_C CH–C–), 6.17 (m, 2H, H_2_C C*H*–C–), 5.85 (broad s, 2H, –C–N*H*–CH_2_–), 5.61 (dd, 2H, *H*_2_C CH–C–), 4.20 (m, 4H, – H_2_C–C*H*_2_–O–), 3.65 (m, 1720H, –O–C*H*_2_–CH_2_–), 3.34 (m, 4H, –CH_2_–C*H*_2_–NH–). The end-group functionalization of the PEUDAm polymers used in this work were greater than 80%.

### Hydrogel coating of radiofrequency ablation catheters

PEUDAm was dissolved in DI water, along with 1 wt% NAGA and photoinitiator Irgacure-2959 at a final concentration of 0.1 wt/vol%, forming the hydrogel precursor solutions. Three compositions were fabricated, each with varying molecular weight and concentration of PEUDAm: 1) 10 wt% 20 kDa, 2) 20 wt% 10 kDa, and 3) 30 wt% 3.4 kDa. For each coating, the metal catheter tip (Biosense Webster EZ Steer with 8mm electrode) was cleaned with acetone, ethanol, and DI water, then dried at 115 °C for 1 hour. The catheter was placed in an oxygen plasma treatment chamber at 18 W and 200 mTorr for 9 minutes, then immediately placed in a solution of 10 vol% 3-(trimethoxysilyl) propyl methacrylate (TMSPMA) in ethanol. After 1 hour, the catheter trip was rinsed in ethanol and dried under nitrogen. Clean glass mold (inner diameter 4 mm) rinsed in Sigmacote was dried at 115 °C for 1 hour prior to filling with the hydrogel precursor solution. The catheter tip was then submerged in the solution and the system placed on a UV transilluminator (UVP, 25 W, 365 nm) for 4 minutes. The catheter tip was carefully removed from the glass mold, and the hydrogel coating was rinsed with deionized water prior to drying overnight at atmospheric pressure.

### Effect of drying, sterilization, and swelling on hydrogel coating integrity

After drying overnight at atmospheric pressure, the dried coated catheter was sterilized with ethylene oxide (EtO). Stereoscope images were taken before and after the sterilization process, as well as after swelling the hydrogel coating to equilibrium in deionized water, to identify any delamination or breakage of the hydrogel (n=3 for each composition, as stated above).

### Simulated deployment-induced damage testing

To assess mechanical durability, each coated catheter was advanced through a 14 Fr Gore DrySeal introducer sheath after equilibrium swelling and then retracted to simulate the most mechanically demanding phase of deployment (n=3 for each composition). Stereoscope images were taken before and after passage through the introducer sheath to examine the coating for damage or delamination. Electrical durability of the hydrogel coating was then assessed; each equilibrated coated catheter was connected to a Biosense Webster SmartAblate system and submerged in a glass tank filled with saline circulating at 37 °C. Radiofrequency ablation cycles were then simulated by performing 50 cycles of applied 30 W for 50 cycles (30 seconds on, 30 seconds off) to mimic a clinical procedure (n=3 for each composition). Stereoscope images were taken before and after ablation cycling to inspect the hydrogel for delamination and microbubble formation.

### Characterization of hydrogel stiffness

To investigate the effect of coating stiffness, three hydrogel compositions were identified to achieve stiffnesses above, below, and matching the stiffness of native atrial tissue.^32^ PEUDAm hydrogel discs were prepared by dissolving PEUDAm and N-acryloyl glycinamide (NAGA) in DI water. Three compositions were fabricated (n=3), each with varying molecular weight and concentration of PEUDAm: 1) 10 wt% 20 kDa, 2) 20 wt% 10 kDa, and 3) 30 wt% 3.4 kDa. A photoinitiator solution of Irgacure-2959 (10 wt/vol% solution in 70 vol% ethanol) was added to these precursor solutions at a final concentration of 0.1 wt/vol%. Hydrogel slabs were fabricated by pipetting the precursor solutions between glass plates (1.5 mm gap) and curing on a UV transilluminator (UVP, 25 W, 365 nm) for 6 minutes on each side. Hydrogel discs (D = 8 mm) were punched from hydrogel sheets and placed into a 12 well plate (n = 3 for each composition) and swollen to equilibrium in deionized water over 24 hours with three solution exchanges. The modulus of the hydrogel discs was then measured at 2% strain using a TA RSA III dynamic mechanical analyzer in compression mode for a cylindrical geometry.

### Effect of hydrogel coating on contact area with atrial tissue mimic

To fabricate an atrial tissue mimic, 3D-printed molds were filled with a hydrogel precursor solution composed of 20 wt% 10kDa polyethylene glycol diacrylate (PEGDA) aqueous solution with 0.1 wt/vol% Irgacure-2959 and orange food dye, then cured via UV transillumination. This composition was selected because the stiffness (226.3 ± 5.9 kPa) matched reported atrial tissue stiffness ranges.^32^ After preparation of the tissue mimic, the hydrogel coated catheter tip was submerged in DI water (mixed with blue food dye for enhanced visualization) until equilibrium swelling was reached. The catheter was placed in scaffolding such that it contacted the tissue mimic vertically, and a 20 g weight was attached to the catheter to standardize contact force. Images of the cross-sectional contact point between the hydrogel coating and the tissue mimic at various trabeculation geometries were captured. To study the effect of coating thickness on tissue contact, the study was performed with the hydrogel composition held constant at 10% 20kDa PEUDAm. The inner diameter of an acrylic mold was varied from 4 mm to 6 mm, and the bare metal catheter tip was used as a control (n=3 for each mold diameter). This study was repeated with the three hydrogel compositions stated above and the bare metal catheter as a control (n=3 for each composition). Contact area was calculated with Equation (1): *A* = π ∗ (*D*/2)^2^, where *D* is the diameter of the contact interface between the probe and the tissue mimic. All images used for analysis are uploaded to the Texas Data Repository.

### Characterization of lesion formation in an ex vivo model

To examine ex vivo ablation lesions from each coating composition, 30 second radiofrequency ablations at 30 W and 50 W, as described above, took place on ventricular free wall tissue from an excised porcine heart (n=3 for each stiffness). Steam pop occurrence was recorded throughout the study. After the ablations were complete, cross-sectional slices of the lesions were taken and stained with 2,3,5-triphenyltetrazolium chloride (TTC), after which images were taken with a Canon EOS 4000D camera to quantify lesion width and depth using ImageJ. The bare metal catheter was used as a control in all cases.

### Characterization of lesion formation in a tissue mimic model

To facilitate more rapid iterative testing of ablation lesion formation, we developed a tissue mimic model. To fabricate the atrial tissue mimic, one well of a 6-well cell culture plate was filled with a hydrogel precursor solution composed of 20 wt% 10kDa PEGDA, 0.1 wt/vol% Irgacure-2959 and 0.3 wt/vol% thermochromic dye (SFXC) then cured via UV transillumination and swelled to equilibrium in 0.9% saline. This hydrogel was engineered to match the mechanical and electrical properties of native atrial tissue and incorporates a thermoresponsive color change (black-to-white at 60 °C) aligned with the temperature threshold for irreversible tissue injury during ablation.^37–39^ The ablation study was repeated with 30 second radiofrequency ablations at 30 W and then the tissue mimic was removed from the saline tank and cross-sectional slices of the lesions were imaged to quantify lesion width and depth as described above.

### Statistical analysis

The data for all measurements are displayed as mean ± standard deviation. An analysis of variation (ANOVA) comparison utilizing Tukey’s multiple comparison test was used to analyze the significance of data among multiple compositions. All tests were carried out at a 95% confidence interval (p < 0.05): ∗ (p < 0.05), ∗∗ (p < 0.001), ∗∗∗ (p < 0.0002), ∗∗∗∗ (p < 0.0001).

## Acknowledgements

This work was supported by the National Institutes of Health, grant number R01 HL162741 (E.C.H., M.R.). Any opinions, findings, and conclusions or recommendations expressed in this material are those of the author(s) and do not necessarily reflect the views of the National Institutes of Health. The renderings in the graphical abstract and Figures 1, 2, 5, and 8 were created in BioRender.

## Table of Contents Figure

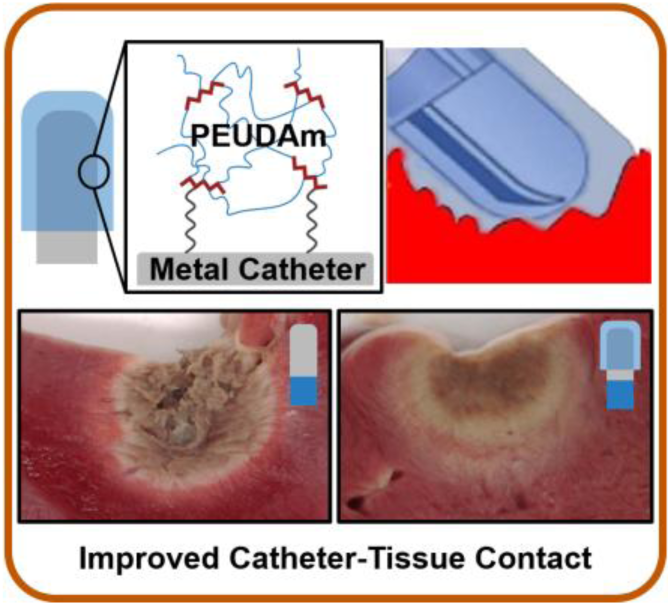
**TOC.** We established a coating method to apply a soft, conductive hydrogel coating to radiofrequency ablation catheters that was mechanically and electrically durable. This conformable coating increased contact area with atrial tissue, increased lesion homogeneity and prevented tissue cavitation due to steam pops.

