## Supplemental Information for "Design of a Conductive Hydrogel Coating to Improve Catheter–Tissue Coupling in Radiofrequency Ablation"

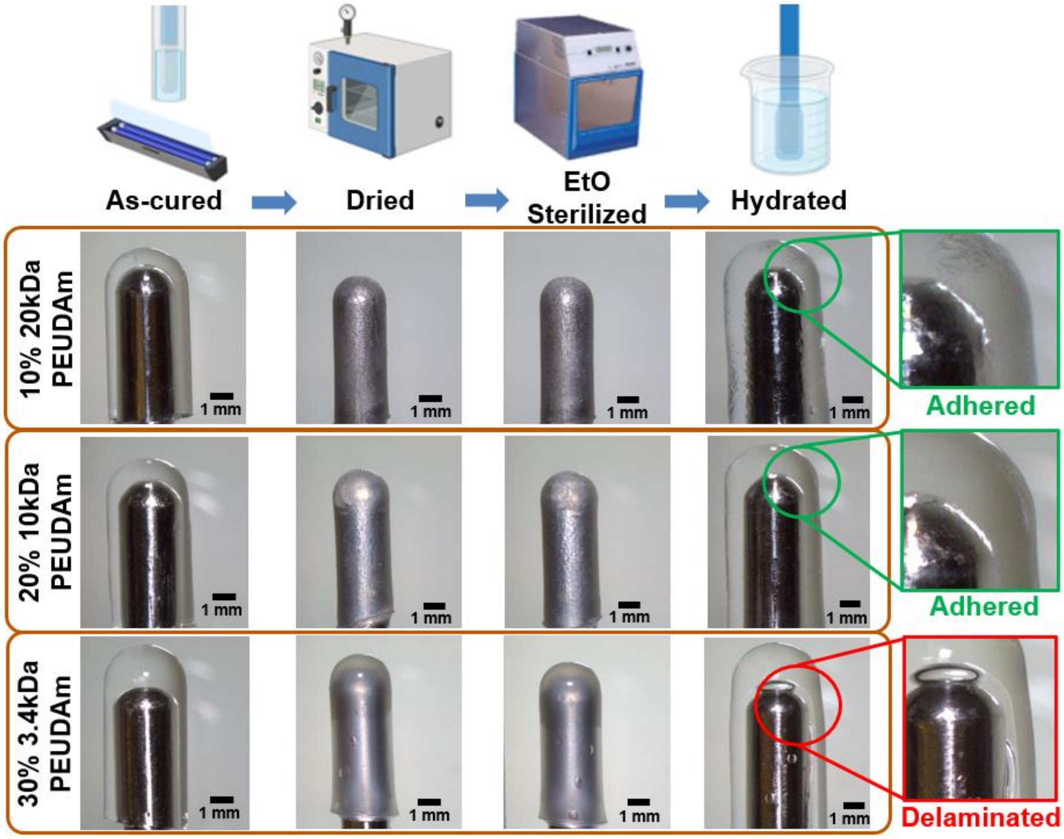


**Figure S1. Hydrogel coating physical integrity after sterilization protocol.** Stereoscope images of each coating composition before and after drying, Ethylene Oxide (EtO) sterilization, and swelling to equilibrium in DI water. n = 3 for each composition.

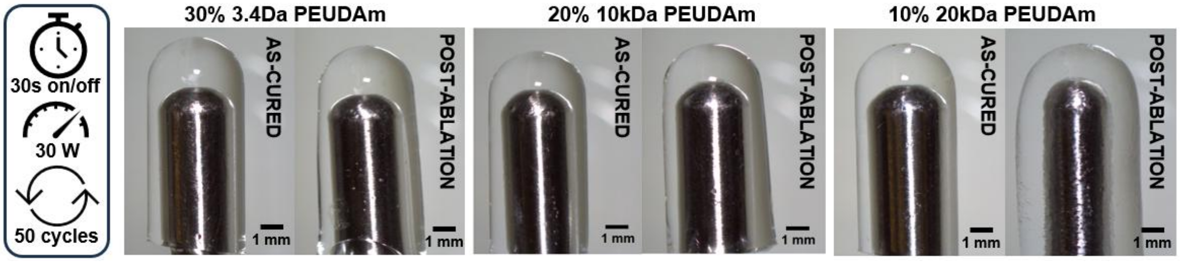


**Figure S2. Effect of radiofrequency ablation cycling on hydrogel coating integrity.** Stereoscope images of the coating before and after deployment durability testing. Each coating undergoes 50 ablation cycles (30 seconds on, 30 seconds off) at 30 W. n = 3 for each composition.

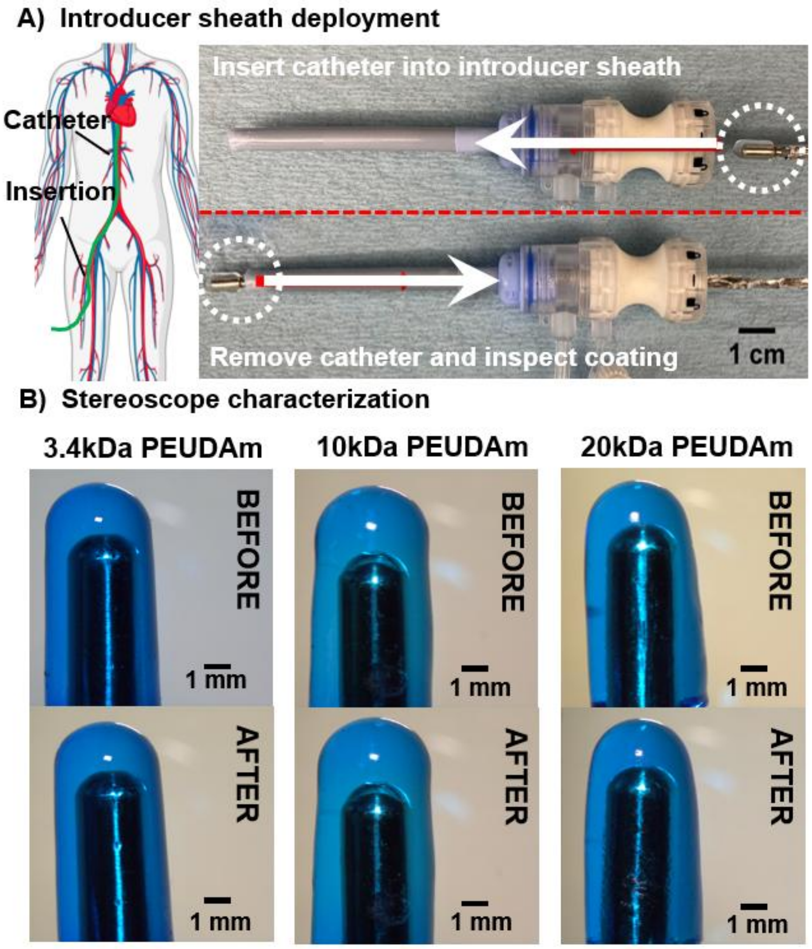


**Figure S3. Assessment of damage after simulated deployment.** a) Each coating undergoes one pass both ways through an introducer sheath. b) Stereoscope images of the coating before and after deployment durability testing. n = 3 for each composition. Figure created with BioRender.com.

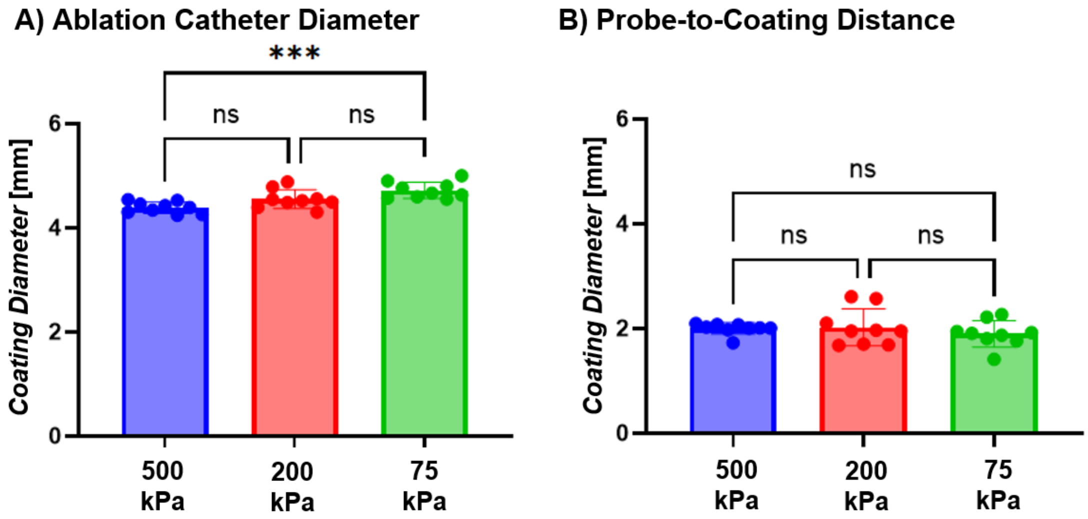


**Figure S4.** Hydrogel coating uniformity based on a) coating diameters and b) distance from probe to tip. Data for all comparisons represent individual data points, means, and standard deviations of each specimen (n = 9 for each composition, 3 compositions). Significant differences between groups are marked as follows: ∗∗∗ ($p < 0.0002$).

**Table S1.** Hydrogel coating uniformity based on coating diameters and distance from probe to tip compared to contact area with tissue mimic (n = 9 for each composition, 3 compositions).

| | Bare Metal | 500 kPa | 200 kPa | 75 kPa |
|---|---|---|---|---|
| Ablation catheter diameter [mm] | 2.32 ± 0.01 | 4.39 ± 0.11 | 4.55 ± 0.18 | 4.72 ± 0.16 |
| Distance from probe tip to outer edge of hydrogel [mm] | - | 2.00 ± 0.11 | 2.02 ± 0.35 | 1.90 ± 0.25 |
| Contact Area [$mm^2$] | 2.88 ± 0.47 | 5.36 ± 0.68 | 8.27 ± 1.15 | 15.21 ± 1.28 |

**Table S2.** Mold-formed coating diameters (n = 3).

| | 4 mm | 6 mm |
|---|---|---|
| Swelled coating diameter [mm] | 5.21 ± 0.13 | 6.16 ± 0.25 |
